# NetMedPy: A Python package for Large-Scale Network Medicine Screening

**DOI:** 10.1101/2024.09.05.611537

**Authors:** Andrés Aldana, Michael Sebek, Gordana Ispirova, Rodrigo Dorantes-Gilardi, Albert-László Barabási, Joseph Loscalzo, Giulia Menichetti

## Abstract

**Summary:** Network medicine leverages the quantification of information flow within sub-cellular networks to elucidate disease etiology and comorbidity, as well as to predict drug efficacy and identify potential therapeutic targets. However, current Network Medicine toolsets often lack computationally efficient data processing pipelines that support diverse scoring functions, network distance metrics, and null models. These limitations hamper their application in large-scale molecular screening, hypothesis testing, and ensemble modeling. To address these challenges, we introduce NetMedPy, a highly efficient and versatile computational package designed for comprehensive Network Medicine analyses.

**Availability:** NetMedPy is an open-source Python package under an MIT license. Source code, documentation, and installation instructions can be downloaded from https://github.com/menicgiulia/NetMedPy and https://pypi.org/project/NetMedPy. The package can run on any standard desktop computer or computing cluster.

## 1 Introduction

Network medicine is a post-genomic discipline that harnesses network science principles to analyze the complex interactions within biological systems, viewing diseases as localized disruptions in networks of genes, proteins, and other molecular entities (Barabási et al., 2011). By integrating comprehensive biological networks, such as the interactome or protein-protein interaction network (PPI), with databases of disease-associated genes (GDA) and ligand-protein interactions, Network Medicine has: 1) successfully identified functional pathways linked to specific phenotypes and diseases (Sharma et al., 2015); 2) pinpointed potential drug targets, highlighting opportunities for both drug repurposing (Cheng et al., 2018, Patten et al., 2022) and effective drug combinations (Cheng et al., 2019). Additionally, this framework has been extended beyond pharmaceuticals to identify food-derived small molecules that impact specific therapeutic areas(do Valle et al., 2021, Nasirian and Menichetti, 2023).

The structure of the biological network plays an essential role in the system’s ability to efficiently propagate signals and withstand random failures. Consequently, most analyses in Network Medicine focus on quantifying the efficiency of the communication between different regions of the interactome. For example, proteins involved in similar therapeutic areas or disease modules are expected to create a cohesive functional subgraph of proteins effectively communicating and influencing each other. In turn, diseases with high pathobiological similarity typically reside in overlapping neighborhoods of the interactome as measured by the *separation* score (Menche et al., 2015, Supplementary Information SI). Similarly, areas of the interactome perturbed by a drug should be close to its protein targets as quantified by the *proximity* score (Guney et al., 2016, Supplementary Information SI).

The speed and reliability of signaling are most commonly quantified through shortest-path metrics with expectations set by uniform or degree-preserving null models, highlighting biological properties not solely determined by link density or degree distribution (Supplementary Information SII). However, biological information does not always travel along geodesic paths, in part because of differences in the flow of information across links. Therefore, a comprehensive assessment of proximity and separation must consider additional metrics of diffusion and communicability. Given the complexity of these combinatorial settings, efficient algorithms are crucial for exhaustive screenings of disease atlases and molecular libraries.

Despite the importance of network measures, most Network Medicine packages focus on the curation of the interactome (Helmy et al., 2022, de Carvalho, 2023) or the curation of GDAs (de Weerd et al., 2022, Ben Guebila et al., 2023). Existing packages that calculate proximity and separation have not advanced beyond their initial introduction (Wang et al., 2022, Maier et al., 2024). As a result, these tools remain highly inefficient for large-scale screening and rely exclusively on shortest-path metrics and limited sample size for hypothesis testing. Here, we introduce NetMedPy, an intuitive Python package for Network Medicine designed to quantify network localization, calculate proximity and separation between biological entities, and conduct screenings involving a large number of diseases and drugs efficiently. NetMedPy provides users with four default metrics and null models with automated statistical analyses. Optimized for high performance in large-scale studies, NetMedPy enhances the robustness and scalability of Network Medicine research, facilitating the discovery of mechanisms of action and prioritizing hypotheses for experimental validation.

## 2 NetMedPy

The workflow of NetMedPy, as illustrated in Figure 1A, involves: 1) loading the interactome, 2) computing and storing the distance matrix induced by a selected metric, 3) loading the desired GDAs and drug targets, and 4) calculating the selected scoring functions (proximity, separation) with the null models of choice. The pipeline output can be further used in downstream analyses. NetMedPy supports weighted and unweighted networks through a Graph object in NetworkX, a widely used library for network analysis. GDAs are entered using a dictionary format, where keys represent disease names and values are lists of associated genes. A similar approach is used for drug targets. The results are then returned in dictionaries, detailing the statistical analysis performed for proximity and separation. For large-scale screening studies, the output is stored in tabular form using Pandas DataFrames.

**Figure 1:**
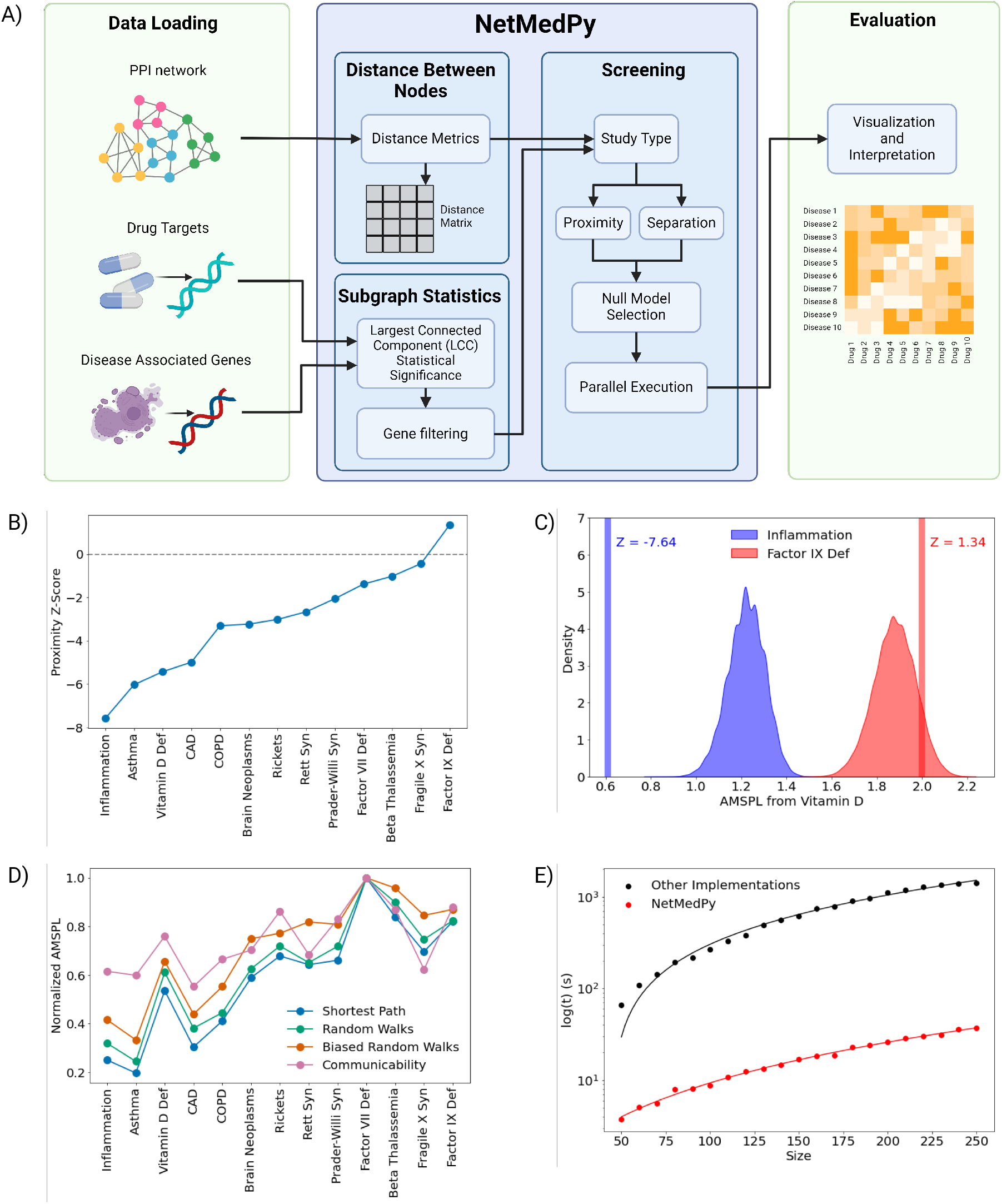
Overview and application of NetMedPy. **>A)** Diagram of the NetMedPy pipeline. Users first load an interaction network, drug targets, and GDAs. NetMedPy calculates the distance matrix induced by the chosen metric for all nodes in the network. Then users set options for subgraph statistics, study type (e.g., proximity, separation), null model, and execution parameters. Visualization and interpretation are performed outside of NetMedPy. **B)** Proximity between Vitamin D’s targets and various diseases. A large negative Z-score indicates a statistically significant closeness between Vitamin D and the disease, while Z-scores close to zero are no different from random. **C)** AMSPL distribution and proximity Z-scores for Vitamin D to Inflammation and Factor IX Deficiency, comparing Vitamin D’s targets to disease genes (vertical lines) and degree-preserving log-binned null models (density plots). Inflammation shows a significantly smaller AMSPL. **D)** Normalized AMSPL using different distance metrics: Shortest Path (blue), Random Walks (green), Biased Random Walks (orange), and Communicability (pink). **E)** NetMedPy execution time (red) versus Proximity implementations found in other packages (black; PMC11223884, PMC4740350, PMC9374494) for increasing gene set sizes. Dots represent time measurements, and straight lines indicate quadratic functions fitted to the data. In each experiment, the proximity Z-Score was calculated using one hundred random samples for illustration purposes. All calculations were performed with a 10-core Intel i9-12900H processor and 32 GB of RAM.

NetMedPy offers a comprehensive suite of metrics, including shortest paths (Menche et al., 2015), random walks (Masuda et al., 2017), biased random walks (Erten et al., 2011), communicability (Estrada and Hatano, 2008), and user-defined options (Supplementary Information SIII). This wide range of metrics allows researchers to tailor their analysis to the specific requirements of various biological questions. The ability to define custom metrics further empowers researchers to develop specialized approaches for their unique research needs. By applying ensemble learning techniques, researchers can also combine the strengths of diverse metrics, enhancing the reliability and depth of their conclusions. This integrated approach can help prioritize experimental tests, improving cost-efficiency and reducing the time and effort required for validation.

NetMedPy provides primary functions for the analysis of disease modules, proximity, separation, and largescale screening studies, including:

- Modules: Given an interactome and a set of nodes, *A*, NetMedPy extracts the largest connected compo-nent (LCC) or subgraph formed by set *A* and calculates the statistical significance of the LCC size (Supplementary Information SI.I).
- Proximity: The original proximity measure *P* (*A, B*) between node sets *A* and *B* is asymmetric, meaning that *P* (*A, B*) *≠ P* (*B, A*). NetMedPy addresses this property by offering both an asymmetric and symmetric proximity *P*_*s*_(*A, B*) Z-score (Supplementary Information SI.II).
- Separation: NetMedPy calculates separation (Supplementary Information SI.II) and its statistical significance, expressed by Z-Score and P-value.
- Screening: NetMedPy incorporates a screening function to calculate network measures between sets of diseases and drugs. The function runs in parallel, enhancing the computational efficiency of multi-core processing capabilities.

Network Medicine leverages null models that generate random samples as benchmarks. By comparing observed network measures against these null hypotheses, researchers can confidently assert the non-randomness of their findings, thereby substantiating the biological relevance of the observed relationships. NetMedPy enhances the robustness of this statistical analysis by incorporating various null models: Perfect Degree Match, Logarithmic Binning, Strength Binning, Uniform Distribution, and user-provided models (Supplementary Information SII). Each null model selects random node sets differently, allowing researchers to account for diverse network properties and biases that might influence the analysis.

## 3 Case Study with Vitamin D

To showcase NetMedPy, we evaluated the role of Vitamin D for an array of 13 disease phenotypes and endophenotypes, selected based on the strength of experimental evidence supporting Vitamin D as a treatment. These categories include strong support (Inflammation, Asthma, Coronary Artery Disease (CAD), Vitamin D Deficiency, Chronic Obstructive Pulmonary Disease (COPD), Rickets), medium support (Brain Neoplasms, Rett Syndrome), and low support (Prader-Willi Syndrome, Factor VII Deficiency, Beta Thalassemia, Fragile X Syndrome, Factor IX Deficiency). We curated Vitamin D’s drug-target data (Piras et al., 2024) and the GDAs of each therapeutic area with and without experimental evidence of Vitamin D modulation (Supplementary Information SIV). Vitamin D is known to 1) reduce the activity of pro-inflammatory cells, 2) regulate blood pressure, and 3) reduce proliferation and boost apoptosis of cancer cells by regulating gene expression via Vitamin D receptors. Leveraging an interactome that integrates the protein-protein interactions reported in (Luck et al., 2020), (Huttlin et al. [2021]), and (Maron et al., 2021) we calculate the proximity between Vitamin D’s targets and each GDA set (Figure 1B).

Our findings reveal that the observed average minimum shortest path length (AMSPL) between Vitamin D and inflammation is significantly smaller than expected when considering node sets of the same size and comparable degree (Z-Score = -7.64), confirming that Vitamin D influences inflammatory processes. Conversely, Factor IX Deficiency, a Mendelian disorder, is more distant from Vitamin D’s targets than expected by chance (Z-Score = 1.34), providing a reasonable negative result (Figure 1B-C). When evaluating the proximity values between Vitamin D and all selected phenotypes, we find that inflammation and related diseases such as asthma show the closest proximity to Vitamin D. This result stands in contrast to diseases with no known association to Vitamin D (e.g., Prader-Willi Syndrome, Factor VII deficiency, Beta Thalassemia, Fragile X Syndrome, Factor IX Deficiency), aligning with existing literature (Figure 1B). Finally, the AMSPL-equivalents for four different metrics display a robust ranking of the results under different notions of distance (Figure 1D and Supplementary Figure S1).

## 4 NetMedPy Performance Evaluation and Comparison

Quantifying the statistical significance of network measures such as proximity and separation in large networks is computationally intensive, as it necessitates comparing selected node sets with randomly generated ones to obtain Z-scores and empirical p-values (Supplementary Information SI-SII). NetMedPy leverages parallelism and pre-calculated distances between all pairs of nodes to enhance performance. This optimization allows distances to be computed once and reused multiple times, significantly improving efficiency and facilitating large-scale screening studies. Figure 1E illustrates the execution time of NetMedPy for calculating proximity between random node sets of increasing size. Our findings show that NetMedPy completes this task faster than the regular proximity implementation, found in different Network Medicine packages(Wang et al., 2022, Maier et al., 2024, Patten et al., 2022), even accounting for the time required to pre-calculate the distances. Consequently, as the number of disease genes and drug-disease pairs increases, NetMedPy demonstrates a substantial performance improvement.

## 5 Discussion

We developed NetMedPy, a user-friendly Python package designed to optimize tools for Network Medicine applications. Tailored for high-performance computing, NetMedPy efficiently handles large-scale data, making it ideal for studies involving drug screening, drug repurposing, and comorbidity identification. The package offers functionalities for extracting the LCC and calculating proximity and separation between node sets, with options for both symmetric and asymmetric measures. Additionally, it supports various null models to validate the statistical significance of network metrics, ensuring robust analytical outcomes. NetMedPy is compatible with both weighted and unweighted networks, and results are conveniently generated in dictionaries or Pandas DataFrames for detailed analyses.

The versatility of NetMedPy, with its support for multiple distance metrics and null models, extends its value to numerous scientific fields that utilize networks. For example, it can enhance social network analysis by investigating social interactions and information dissemination. In epidemiology, NetMedPy can analyze disease spread and the effectiveness of health interventions within interconnected populations.

In conclusion, NetMedPy is a valuable tool for researchers, enabling them to uncover new insights and address complex problems with efficient network analysis techniques.

## Acknowledgments

We thank Bnaya Gross for their help in testing the package during development. We thank Ruisheng Wang for the process to curate disease-gene associations and Andrea Piras for the collection of drug-target associations.

## Author Contributions

A.A. contributed the formal analysis, validation, and visualization. A.A., M.S., G.I., and R.D-G. contributed to the methodology and the software. A.A., M.S., G.I., J.L., A.-L.B., and G.M. contributed to the writing and editing.

G.M. conceptualized, administered, and funded the project.

