## Supplementary Information for "NetMedPy: A Python package for Large-Scale Network Medicine Screening"

### NetMedPy: Supplementary Information

**SIV. Selection of Disease Genes** **8**

**SV. NetMedPy Installation** **9**

**SVI. Supplementary Figures** **10**

Agreement between metrics for the AMSPL . . . . . . . . . . 10

### SI. Network Medicine Background

#### SI.I Disease Modules

Different studies have shown that genes associated with a particular disease tend to cluster within the same network neighborhood, frequently forming a connected sub-network within the interactome[1,9,11]. The largest connected component (LCC) formed by these disease genes contains important information about the disease localization within the interactome, as well as the main interactions between the disease genes[11]. Studying the LCC allows researchers to efficiently narrow down the vast number of potential biomarkers to those most likely to be clinically relevant in drug discovery and repurposing efforts.

Calculating the statistical significance of the LCC size in Network Medicine is the standard framework for assessing the relevance of disease gene localization. For any set of nodes in the interactome, the statistical significance of the LCC size is determined by the Z-Score of the observed size $l$:

$$Z = \frac{l - \mu_l}{\sigma_l}, \tag{1}$$

where $\mu_l$ is the expected LCC size for nodes chosen randomly, and $\sigma_l$ is the LCC standard deviation across the randomized ensemble. Additionally, the p-value associated to $l$ is calculated as the probability of observing an LCC of size $l$ or superior in a random sample of nodes. These two indicators are important in their ability to validate the biological relevance of the LCC, ensuring that the connections and interactions within the module are not random artifacts but likely reflect accurate biological processes and pathways implicated in the disease.

#### SI.II Proximity

Given $X$, a set of disease genes and $Y$, a set of drug targets, we define $d_c(X,Y)$ as the average shortest path length between the drug's targets and the nearest disease node[5]:

$$d_c(X,Y) = \frac{1}{|Y|} \sum_{y \in Y} \min_{x \in X} d(x,y), \tag{2}$$

where $d(x,y)$ is the length of the shortest path between node $x$ and $y$. For each drug-disease pair, proximity assesses whether the drug targets are closer to the disease nodes than expected if both sets were randomly chosen across the network. The statistical significance of the distance $d_c$ is determined by calculating the Z-score of the observed drug-disease distance compared with the respective random expectation:

$$Z = \frac{d_c - \mu}{\sigma}. \tag{3}$$

Large negative scores indicate that the drug targets are closer to disease nodes than random, while large positive scores indicate the drug targets are farther away from disease nodes than expected. The p-value of $d_c$ is calculated as the probability of observing a distance of length $d_c$ or inferior in random samples of drug targets and disease nodes. Note that Eq. (2) is not symmetric, since $d_c(X,Y) \neq d_c(Y,X)$.

To broaden the applicability of proximity beyond its original use in drug discovery, we propose a straightforward symmetric proximity measure by replacing $d_c$ in Eq. (2) with the average minimum shortest path length in both directions:

$$d_s(X,Y) = \frac{d_c(X,Y) + d_c(Y,X)}{2}. \tag{4}$$

Then, $\mu$ and $\sigma$ in Eq. (3) are calculated using a null model that considers the definition of $d_s$ in Eq. (4).

#### SI.III Separation

Similar to proximity, separation was introduced as a relative measure of distance between node sets within a network, with its primary application being the identification of disease comorbidities[8]. Indeed, if two disease modules overlap, perturbations leading to one disease are most likely to propagate to the other disease module, disrupting pathways and biological processes involved in both diseases and resulting in shared abnormalities and phenotypes. The separation between disease $A$ and disease $B$ is defined as

$$S_{AB} = d_{AB} - \frac{d_{AA} + d_{BB}}{2}, \tag{5}$$

where $d_{AB}$ represents the average of the shortest path length between all genes of disease $A$ and all genes of disease $B$. Hence, $S_{AB}$ compares the shortest distance between genes of diseases $A$ and $B$ to the within-disease-module distance $d_{AA}$ and $d_{BB}$. Negative separation values indicate that the average shortest path distance between the modules is smaller than the "average size" of the two modules, suggesting the presence of shared mechanisms or risk factors.

### SII. Null Models in NetMedPy

NetMedPy offers the following null models to estimate the significance of the LCC, proximity, and separation:

- **Perfect Degree Match.** This model selects a random sample replicating the original node-set's degree distribution. A node's degree, indicating its number of connections, is a highly predictive network property. By preserving this distribution in the sample, the null model ensures that the random set is structurally similar to the original, allowing to search for patterns not solely driven by the degree.
- **Logarithmic Binning.** This model involves categorizing the degrees of all nodes

within the network into logarithmically sized bins, with each bin containing a fixed number of nodes. Samples are then drawn by matching the degree of the original nodes to those within the corresponding bins. This method allows more variability for high degree nodes, providing a more generalized yet representative sample for comparison.

- **Strength Binning.** Analogous to logarithmic binning, strength binning uses the strength of the nodes—typically a weighted sum of their connections—instead of their degrees. This model is particularly useful in weighted networks where the strength of the nodes might provide a more accurate reflection of their functional significance.

- **Uniform Distribution.** This model randomly selects nodes from the entire network, disregarding their degree or strength. It represents an entirely random sampling approach, providing a baseline for evaluating the extent to which the original node set's structural or functional features influence the observed metrics.

- **Custom Null Model.** This feature allows users to specify their null model, enabling the application of unique or novel statistical frameworks adapted to specific research questions or datasets. It provides methodological flexibility and can be particularly valuable in cases where standard null models do not adequately address the particularities of the data.

### SIII. Information passing metrics

Proximity and separation rely on the definition of a "distance" between nodes, measured as the length of their shortest path, to quantify how information traverses the interaction network.

Beyond the shortest path, NetMedPy includes three additional methodologies to quantify information flow within the network: random walks with restart (RWR), biased random walks with restart (BRWR), and communicability. Each method calculates an association score $\phi(a,b)$, reflecting how effectively information can be transmitted between node $a$ and $b$. Large $\phi(a,b)$ scores in any of these measures suggest a strong potential for interaction or functional relationship between nodes. Consequently, we redefined the distance between nodes based on $\phi(a,b)$, ensuring that higher scores correspond to shorter distances within the network.

#### SIII.I Random Walks with Restart

Random Walks with Restart (RWR) simulate the random movement of molecules or signals through a network, allowing the walker to return to a starting node with a certain probability at each step. This approach captures the stochastic nature of biological processes while emphasizing the local context of the starting node. The restart mechanism ensures that the walker frequently returns to the initial node, preventing it from drifting too far and providing a more accurate reflection of node importance within the vicinity. This makes RWR valuable for understanding biological networks' local and global properties and identifying key regulatory nodes and potential therapeutic targets[6].

Let $A$ be the adjacency matrix of the network, $D$ be the diagonal degree matrix where $D_{ii} = \sum_j A_{ij}$, $P$ be the transition probability matrix given by $P = AD^{-1}$, and $\alpha$ be the restart probability at node $a$. The steady-state probability distribution $r_a$ of a random walk restarting at node $a$ is defined as[7]:

$$r_a = \alpha(I - (1-\alpha)P)^{-1}p, \tag{6}$$

where $p$ is the restart distribution vector, typically a unit vector with 1 at the starting node and 0s elsewhere. Specifically, each element $r_{ai}$ of the vector $r_a$ corresponds to the steady-state probability of being at node $i$ after a large number of steps in the random walk process, considering the possibility of restarting at the initial node $a$ with probability $\alpha$. The association score $\phi_{RWR}(a,b)$ for RWR is simply:

$$\phi_{RWR}(a,b) = r_{ab}. \tag{7}$$

#### SIII.II Biased Random Walks with Restart

Biased Random Walks with Restart (BRWR) enhance RWR analysis by incorporating node-specific biases to mitigate the influence of highly connected nodes[2]. We achieve this by dividing the steady-state probability $r_{ab}$ in Eq. (7) by the degree of node $b$, $k_b$. Then, the association score $\phi_{BRWR}(a,b)$ for BRWR is calculated as:

$$\phi_{BRWR}(a,b) = \frac{r_{ab}}{k_b}. \tag{8}$$

This technique effectively reduces the bias towards highly connected nodes, providing a more balanced and nuanced understanding of node significance and interaction patterns. BRWR is particularly useful for identifying influential nodes that might not be apparent in the standard RWR.

#### SIII.III Communicability

Communicability considers all possible paths between nodes, weighted by their lengths, thus providing a comprehensive view of their interaction potential. This measure evaluates pathway redundancy by identifying how perturbations can propagate through indirect interactions, offering a deeper understanding of systemic effects in the network[3,4]. The association score $\phi_{COM}(a,b)$, defined as the communicability between nodes $a$ and $b$, follows as:

$$\phi_{COM}(a,b) = \sum_{l=0}^{\infty} \frac{(A^l)_{ab}}{l!} = (e^A)_{ab}, \tag{9}$$

where $l$ is the length of the path.

#### SIII.IV Converting association scores into distances

Since large values of the association score $\phi(a,b)$ are indicative of strong or close interaction between nodes, we define the $\phi$-score based "distance" $d_\phi(a,b)$ by inverting and normalizing $\phi(a,b)$:

$$d_\phi(a,b) = 1 - \frac{log(\phi(a,b)) - m}{M - m}, \tag{10}$$

where $m = min_{a,b}\{log(\phi(a,b))\}$ and $M = max_{a,b}\{log(\phi(a,b))\}$. In this way, the distance $d_\phi(a,b)$ assigns small values to nodes with strong association score $\phi(a,b)$, and values close to 1 as $\phi(a,b)$ decreases.

### SIV. Selection of Vitamin D's drug targets and Disease Genes

Vitamin D's drug targets were extracted using the CPIExtract package, as detailed in[10]. In brief, CPIExtract extracts, filters, and harmonizes raw data from nine databases: BindingDB (BDB), ChEMBL, Comparative Toxicogenomics Database (CTD), DrugBank (DB), DrugCentral (DC), Drug Target Commons (DTC), Open Targets Platform (OTP), PubChem, and STITCH. For Vitamin D, we find a total of 23 targets mapped to the selected version of the human interactome.

Gene-disease associations (GDAs) for each disease were extracted and standardized from DisGeNet, Phenopedia, and Open Targets. To ensure that only associations with strong experimental support were retained, we applied the filtering criteria specified in Table S1.

To balance the number of GDAs from each source, we adjusted these thresholds for Inflammation, COPD (Chronic Obstructive Pulmonary Disease) and CAD (Coronary Artery Disease), according to the values described in Table S2.

| GDA Source | Association Score | Value |
|---|---|---|
| Phenopedia | Total Publications | 1.00 |
| Open Targets | Overall Association Score | 0.10 |
| DisGeNet | Score GDA | 0.01 |

Table S1: Filters applied to produce the GDAs from each source

| Disease | Source | Association Score | Value |
|---|---|---|---|
| Inflammation | Phenopedia | Total publications | 4.00 |
| COPD | DisGeNet | Score GDA | 0.20 |
| CAD | DisGeNet | Score GDA | 0.05 |

Table S2: Filter criteria for specific diseases, to balance the number of disease genes from each source.

### SV. NetMedPy Installation

NetMedPy can be installed locally either from GitHub or using pip. To install from GitHub, navigate to the repository at https://github.com/menicgiulia/NetMedPy and follow the instructions in the README file. Alternatively, you can install the package using pip with the following command:

```
pip install netmedpy
```

The package is designed to run in a Python environment above 3.6 and below 3.12, ensuring full functionality of all features. For detailed installation guidelines and additional information, refer to the documentation available on the GitHub repository.

### SVI. Supplementary Figures

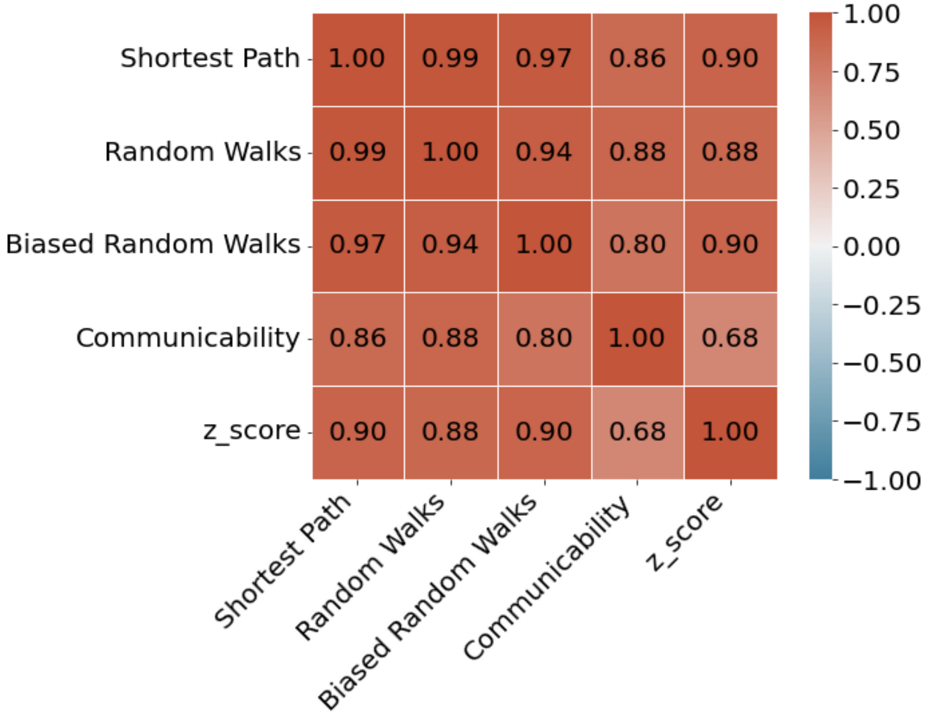

Figure S1: Agreement between the different metrics for the Vitamin D's AMSPL analysis. For the Vitamin D case study, we calculate the Pearson correlation among the AMSPL-equivalent raw values calculated with different metrics, including an additional comparison with the degree-preserving proximity Z-score calculated for the shortest path metric (Figure 1B). The correlation matrix demonstrates strong agreement between the metrics, indicating robustness in the results across different distance notions.
